# Phytochemicals from *Daucus carota* have therapeutic potential against Triple-Negative Breast Cancer PDK1 and PLK1

**DOI:** 10.1101/2024.11.04.621987

**Authors:** Kayode Raheem, Modinat Abayomi, Maryam Oluwatosin, Mary Adewunmi, Ijaz Ali, Muhammad Muddassar

## Abstract

Triple-negative breast cancer (TNBC) is a deadly form of breast cancer that lacks estrogen, progesterone, and HER2 receptors. The development of drugs for TNBC has been challenging due to the lack of specific therapeutic targets. However, recent studies have shown that targeting the ATP active of the PDK-1 and PLK-1 proteins could be potential drug targets for TNBC treatment. New medications for TNBC have considerable adverse effects, highlighting the need for more targeted and effective therapies.

In this study, we employed various computational approaches, including molecular docking, molecular dynamics (MD) simulations, pharmacokinetic studies, binding free energy calculations, principal component analysis (PCA), and alanine scanning analysis, to identify bioactive compounds from *Daucus carota-*extracted natural compounds that can bind to these ATP-binding sites and inhibit the activity of PDK1 and PLK1. Our study revealed that eight compounds showed reasonably good docking scores, binding free energies, and ADMET properties against the PDK1 and PLK1 enzymes. Astragalin and scolimoside showed substantial binding affinity and persistent interactions in the pocket region of the two proteins. Further MD simulation studies for 150 ns also suggested that the compounds were stably bound in the active site with very minor fluctuations. We believe that the identified hits for PDK1 and PLK1 from *Daucus carota* will be effective against TNBC in performing the biological assays.

## 1. Introduction

Triple-negative breast cancer (TNBC) is an aggressive subtype of breast cancer characterized by the absence of estrogen receptor, progesterone receptor, and human epidermal growth factor receptor 2 (HER2) expression ^1^. As a result of the lack of these receptors, hormonal and HER2-targeting therapies are ineffective against TNBC. Therefore, nonspecific chemotherapy is the most prevalent method of treatment ^2^ and has serious side effects, such as skeletal muscle damage, cardiotoxicity, and myelosuppression ^2, 3^. Often, untargeted chemotherapeutic agents must be administered at a lower dose to prevent side effects; however, this results in poorer treatment outcomes and cancer recurrence ^4^. There is a poor 5-year prognosis of 12% for patients who experience TNBC relapse and progress to a metastatic stage ^5^. The development of effective treatment strategies for these highly aggressive TNBC tumors is urgently needed.

As part of their signal transduction functions, protein kinases and other regulatory proteins have evolved stringent switches that control activation, modulating the ATP-binding site (orthosteric site) conformation in response to the appropriate upstream signals ^6^. In breast cancer cells, 3-phosphoinositide-dependent protein kinase 1 (PDK1), a major AGC protein kinase, plays critical roles in multiple cellular processes, including tumorigenesis, growth, survival, metastasis, regulation of the tumor microenvironment (TME), and resistance to drugs, via either the PI3K/AKT-dependent or the PI3K/AKT-independent pathway ^7, 8^. Therapy targeting PDK1 signalling has been proposed to improve BC efficacy or even reverse treatment failure^9^. PDK1 is a cytosolic protein with three ligand binding sites: a substrate binding site, a catalytic ATP binding site, and a PDK1 interacting fragment (PIF) binding site^10^. PIF pockets serve two functions: attracting downstream substrate kinases harboring hydrophobic motifs (HMs) and intrinsically regulating PDK1 activity ^11^. More than 500 protein kinases in the human genome share a relatively conserved ATP binding site ^12, 13^.

Polo-like kinase 1 (PLK1) plays an indispensable role in regulating multiple steps of mitosis, including centrosome maturation, spindle assembly, chromosome separation, and cytokinesis. In most tumors, PLK1 acts as an oncogene during disease progression. Research has shown that inhibiting PLK-1 can decrease tumor initiation and invasion and promote apoptotic cell death, especially in combination with chemotherapy and other targeted therapies^14^. PLK1 has two main domains that can be targeted: a kinase domain (KD) and a polo-box domain (PBD). Many current PLK1 inhibitors are designed to bind to the ATP-binding pocket in the KD or the phosphopeptide-binding site in the PBD^15^. There are more than ten commercially available PLK1 inhibitors, at least 4 of which are ATP-competitive inhibitors that have been evaluated clinically. Recent studies have explored the use of plant-derived compounds as alternative inhibitors of PLK1 and related kinases such as PDK1. Potential motivations include overcoming resistance observed with ATP-competitive inhibitors, taking advantage of the diverse structures and mechanisms of plant metabolites, and benefiting from the longstanding medicinal use of many plant sources. By focusing research on plant derived PLK1 and PDK1 inhibitors, new therapeutic compounds could be discovered.

In this study, we propose the use of Daucus carota (Carrot) phytochemicals as potential inhibitors of PLK1 and PDK1. Daucus carota, commonly known as carrot, has gained significant attention in recent years for its potential as an effective anticancer agent. Several studies have investigated the potent anticancer activity of Daucus carota. In breast cancer cell lines, the pentane fraction and pentane/diethyl ether fraction of Daucus carota oil extract demonstrated remarkable efficacy by arresting the cell cycle, inducing apoptosis ^16^, and suppressing metastasis ^17^. Furthermore, the isolation of 6-methoxymellein from carrots revealed its ability to target breast cancer stem cells through the regulation of NF-κB signalling, offering a promising strategy for combating breast cancer^18^. Daucus carota oil extract exhibited significant in vitro anticancer activity against multiple human cancer cell lines, indicating its broad-spectrum efficacy ^19^. A novel sesquiterpene called β-2-himachalen-6-ol, which was isolated from Lebanese wild carrot, demonstrated noteworthy anticancer properties^20^. Although these studies provide valuable insights into the anticancer potential of Daucus carota, further research is needed to fully elucidate its underlying mechanisms of action. The multifaceted pharmacological properties of Daucus carota, including antioxidant, anti-inflammatory, and hepatoprotective activities, warrant exploration and provide a foundation for developing comprehensive therapeutic strategies ^16, 21^. In this study, we applied a computer-aided drug design approach to discover four potential phytochemical compounds from Daucus carota as novel inhibitors of PLK1 and PDK1 for TNBC treatment.

Using molecular dynamics (MD) simulations, we demonstrated that these lead compounds exhibit persistent binding in the active regions of the PLK1 and PDK1 proteins. The results from our molecular dynamics (MD) analysis revealed positive molecular interactions, including hydrogen bonding and hydrophobic contacts, which help in the binding of phytochemicals. A residue-based free energy decomposition analysis was conducted to determine the binding ability of astragalin, kaempferol 3-O-beta-D-glucoside, quercitrin, scolimoside, luteolin 7-O-(6-malonylglucoside), and thermopsoside to the proteins. A dynamic cross-correlation matrix was utilized to analyse the internal dynamics of the compounds bound to PLK1 and PDK1^22^. This study offers a rational framework for identifying and enhancing natural product inhibitors of PLK1 and PDK1 as novel anticancer drugs^22^.

## 2. Methods and Materials

### 2.1. Compounds Retrieval and Preparation

Phytochemical compounds of Daucus carota were retrieved from Dr. Duke’s Phytochemical and Ethnobotanical Databases ^23^. The structures were prepared using the LigPrep module of the Schrodinger Suite 2020-2. Different ionization and protonation states of the compounds at pH 7 were generated using the Epik module of the Schrodinger Suite. The stereoisomers for each compound with a specified chirality were generated using the OPLS_2005 forcefield. The prepared ligands were then subjected to further analyses and molecular docking studies. The selected ligands with the lowest energy conformations are listed in Table 1S.

### 2.2. Protein preparation

The X-ray crystal structures of the target proteins, polo-like kinase-1 (PLK1 PDB: 3FC2) and protein kinase (PDK1 PDB: 4WA1), with resolutions of 2.45 Å and 1.68 Å, respectively, were retrieved from the Protein Data Bank (http://www.rcsb.org/pdb/) ^24^. The protein structures were processed using the Protein Preparation Wizard module of Schrodinger Maestro 12.0. The prepared protein structures underwent energy minimization and optimization steps, which included removing water molecules and ions, adding hydrogen atoms, assigning bond orders, creating disulfide bonds, adding side chain residues, and adding missing atoms such as missing loops or terminal residues^25^.

### 2.3. Molecular docking

Maestro’s glide module was used for molecular docking studies with the binding pockets of PLK1 and PDK1. The docking grid box was generated around the active site of the proteins with the receptor grid generation panel. The prepared ligands were docked into the respective pocket sites of PLK1 and PDK1 using Glide software. The docking protocol included initial ligand placement, energy minimization, and refinement of the ligand□protein complexes. XP docking (precision tool from Schrodinger software) was performed to assess the lowest binding energy of the docked ligands.

### 2.4. Pharmacokinetic analysis

The optimal ligand□protein conformations from molecular docking were used for further analysis. The pharmacokinetic parameters, such as absorption, distribution, metabolism, and excretion of the ligand□protein complexes, were assessed using SwissADME (http://www.swissadme.ch), a web-based tool for predicting pharmacokinetic properties. The identified ligands were subjected to Lipinski’s Rule of Five, which evaluates drug likeness based on parameters such as molecular weight, lipophilicity, hydrogen bond donor and acceptor count, and the number of rotatable bonds^26, 27^. The toxicity profiles of the selected compounds were evaluated using Data Warrior software^28^. This analysis examined several toxicity endpoints, such as tumorigenicity, reproductive toxicity, irritant effects, and mutagenicity.

### 2.5. Molecular dynamics simulations

The ligands with the lowest energy conformation and the reference compound were subjected to MD simulations to evaluate their conformational flexibility and binding stability using GROMACS (2023) ^29^. The force field parameters used in the MD simulations were implemented using the CHARMM27 force field. The force field parameters for the ligands were generated using the Swissparam (V1.5) server, which is based on the CHARMM General Force Field version 2.0. ^30^. The MMFF was used to determine the topological, atomic, and charge parameters of the ligands. The ligands and protein topologies were merged, solvated, minimized, and equilibrated prior to the MD simulations. The TIP3P water model was used to solvate the system, and counterions were added to neutralize the charge.

Furthermore, Na+ and Cl- ions are required before minimizing the structure of the complex until the maximum force is below 1000 kJ/mol/nm. The NVT and NPT conserved ensembles were generated in the equilibrated system with a simulation time of 100ps for each. The equilibrated system was subjected to production runs of 150 ns with a time step of 2 fs using the Leapfrog algorithm. The analysis of the MD trajectory was performed using XMgrace and the VMD tcl command to analyse various parameters, such as the root mean square deviation, root mean square fluctuation, and radius of gyration^31^.

### 2.6. Binding free energy calculations using MM-GBSA model

Free binding energy/MM-GBSA calculations were performed to evaluate the binding affinity between the ligands and target proteins. The binding free energy was estimated by the difference between the total free energy (ΔG_com_) of the ligand□protein complex and the sum of the free energy of the individual receptors (ΔG_pro_) and ligand (ΔG_lig_) using the equation provided below:

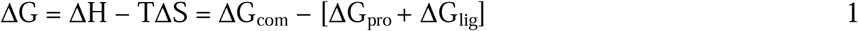

The ΔG for the complex, receptor and ligand can be calculated by the following equation:

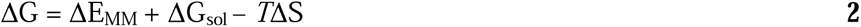

ΔEMM = Molecular Mechanics Energy
ΔGsol = Solvation Free Energy
TΔS = Entropy at a given temperature

### 2.7. Principal component analysis (PCA)

A covariance grid was first evolved using atomic headings obtained from MD bearings to perform the PCA, which were diagonalized to deliver many eigenvalues and eigenvectors. Eigenvectors obtained from the diagonalization of the covariance grid show the direction of advancement in the conformational space of protein regions, while the related eigenvalues reflect the square mean instabilities along the specific eigenvectors. The underlying very few high-regard eigenvectors are particularly useful for illustrating overall protein advancements. The BIO3D group of R was used to determine the dynamic development of the protein structures^32^.

### 2.8. Free energy decomposition analysis

The gmx_MMPBSA of GROMACS was used to calculate the free energy decomposition of the key residues involved in the interaction of ligand and protein. The equation illustrates the interaction between ligand and protein:

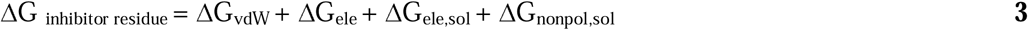

### 2.9. Alanine Scanning Analysis

We employed computational alanine scanning mutagenesis to investigate the contributions of key residues to protein□ligand binding by assessing the impact of replacing their side chains with a smaller, simpler alanine. The binding energies of the mutated complexes were calculated. The total binding free energy of the wild-type and mutated complexes can be calculated with the following equation:

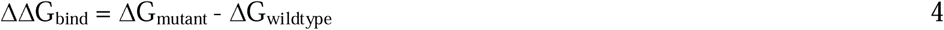

## 3. Results and Discussion

Compared with other breast cancer subtypes, TNBC represents an ongoing treatment challenge due to its lack of targeted therapy, high recurrence rate, and low survival rate. The lack of druggable receptors such as estrogen receptor (ER), progesterone receptor (PR), and human epidermal growth factor receptor 2 (HER2) leaves TNBC patients with limited, targeted therapy options and, consequently, lower survival rates. While recent clinical trials have investigated several new targets, such as P13K/AKT/mTOR, RAS/MAPK, epidermal growth factor receptor (EGFR), and androgen receptor, the treatment of TNBC remains unsolved due to the heterogeneity of the diseases and drug resistance they exhibit.

As evidenced in a previous study, overexpression of PLK1 and PDK1 is associated with tumor development and a poorer prognosis in patients with breast cancer^9, 33, 34^. This revealed that PLK1 and PDK1 could serve as a promising new therapeutic target for ameliorating TNBC progression. However, to the best of our knowledge, directly targeting the ATP-binding sites of these kinases for TNBC treatment using *Daucus carota* as a natural product-based inhibitors has not been extensively explored. In this study, we used phytochemicals extracted from *Daucus carota* to bind to the ATP sites of PLK1 and PDK1. MD simulation analysis, binding free energies analysis, PCA, and alanine scanning analysis revealed two potent phytochemicals, astragalin and scolimoside; that could serve as potential therapeutic agents for treating TNBC.

### 3.1. Molecular docking

The conformers generated were docked against the binding pocket of the PLK1 and PDK1 proteins. The pose with the lowest binding energy was chosen for further analysis (figure 1S), and the potential binding interactions between the ligands and the protein binding sites were evaluated. Eight (8) compounds were selected with high docking scores against the target proteins, as shown in Tables 1 and 2, compared to the reference drug, epirubicin. For the PLK1 protein, astragalin and kaempferol 3-O-Beta-D-glucoside, with a binding affinity of - .9.424 kcal/mol, exhibited the most favourable binding affinity for the protein via hydrogen interactions with the residues ASP 194, LYS 82, CYS 133, and GLU 131. These residues are known to play a crucial role in inhibiting PKL1 activity and have been identified as key binding sites for PLK1 inhibitors in previous literature (Huang et al., 2017). In addition, cyanidin 3,5-diglucosie, chlorogenic acid, scolimoside, thermopsoside, and luteolin-4-O-glucoside have highly promising docking scores for the PLK1 protein, with binding energies ranging from −8.579 to −9.165 kcal/mol. Hydrogen bond interactions, pi stacking, and hydrophobic interactions were observed between the complexes. These interactions underpin the stability, functionality, and integrity of the complexes.

**Table 1:**
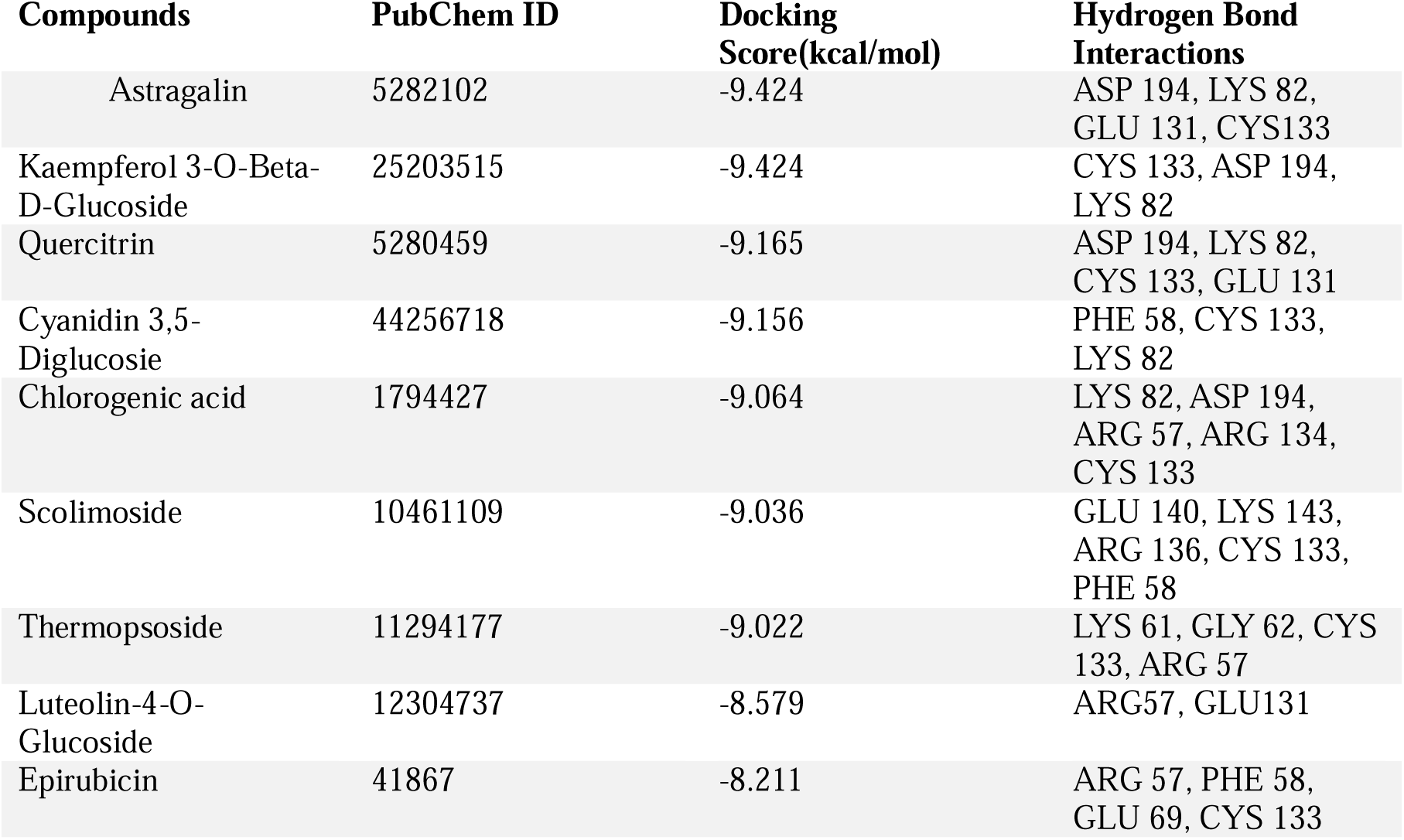
Binding affinity scores of selected ligands against PLK-1 (PDB ID: 3FC2).

| Compounds | PubChem ID | Docking Score(kcal/mol) | Hydrogen Bond Interactions |
| --- | --- | --- | --- |
| Astragalin | 5282102 | -9.424 | ASP 194, LYS 82, GLU 131, CYS133 |
| Kaempferol 3-O-Beta-D-Glucoside | 25203515 | -9.424 | CYS 133, ASP 194, LYS 82 |
| Quercitrin | 5280459 | -9.165 | ASP 194, LYS 82, CYS 133, GLU 131 |
| Cyanidin 3,5-Diglucosie | 44256718 | -9.156 | PHE 58, CYS 133, LYS 82 |
| Chlorogenic acid | 1794427 | -9.064 | LYS 82, ASP 194, ARG 57, ARG 134, CYS 133 |
| Scolimoside | 10461109 | -9.036 | GLU 140, LYS 143, ARG 136, CYS 133, PHE 58 |
| Thermopsoside | 11294177 | -9.022 | LYS 61, GLY 62, CYS 133, ARG 57 |
| Luteolin-4-O-Glucoside | 12304737 | -8.579 | ARG57, GLU131 |
| Epirubicin | 41867 | -8.211 | ARG 57, PHE 58, GLU 69, CYS 133 |

**Table 2:** Binding affinity scores of selected ligands against PDK-1 (PDB ID: 4WA1).

| Compounds | PubChem ID | Docking Score(kcal/mol) | Hydrogen Bond Interaction |
| --- | --- | --- | --- |
| Scolimoside | 10461109 | -10.37 | ASN 210, GLU 209, GLU 90, LYS 169, LYS 163, ALA 162 |
| Luteolin 7-O-(6-Malonylglucoside) | 5281669 | -9.717 | GLU 209, GLU 166, LYS 163, ALA 162, LYS 169, |
| Thermopsoside | 11294177 | -9.122 | GLU 90, LEU 88, ASN 210, GLU 166, LYS 169, ALA 162 |
| Luteolin-7-O-Glucuronide | 13607752 | -9.058 | PHE 93, GLU 209, ALA 162, LYS 163, GLU 166, LYS 169 |
| Apigenin 7-O-Glucoside | 5385553 | -8.905 | GLU 166, ALA 162, ASP 223, GLU 90, LYS 169 |
| Cynaroside | 5280637 | -8.886 | ALA 162, LYS 163, GLU 166, LYS 169, GLU 209, ASN 210 |
| Apigenin-7-O-beta-D-Glucopyranoside | 12304094 | -8.839 | ALA 162, GLU 166, GLU 209, LYS 169 |
| Epirubicin | 41867 | -8.187 | GLU 90, GLU 166, GLU 209, ASP 205, LYS 207, THR 222 |

The results showed that astragalin and kaempferol 3-O-Beta-D-glucoside, cyanidin 3,5-diglucosie, and chlorogenic acid had significant binding affinity to the binding site of the PLK1 protein and their binding pose and 2D interaction, as illustrated in Figure 1A. According to the literature, astragalin has also been reported to exhibit anticancer effects through different mechanisms, including cell cycle arrest and apoptosis induction, in various types of cancer cells^35, 36^.

**Figure 1:**
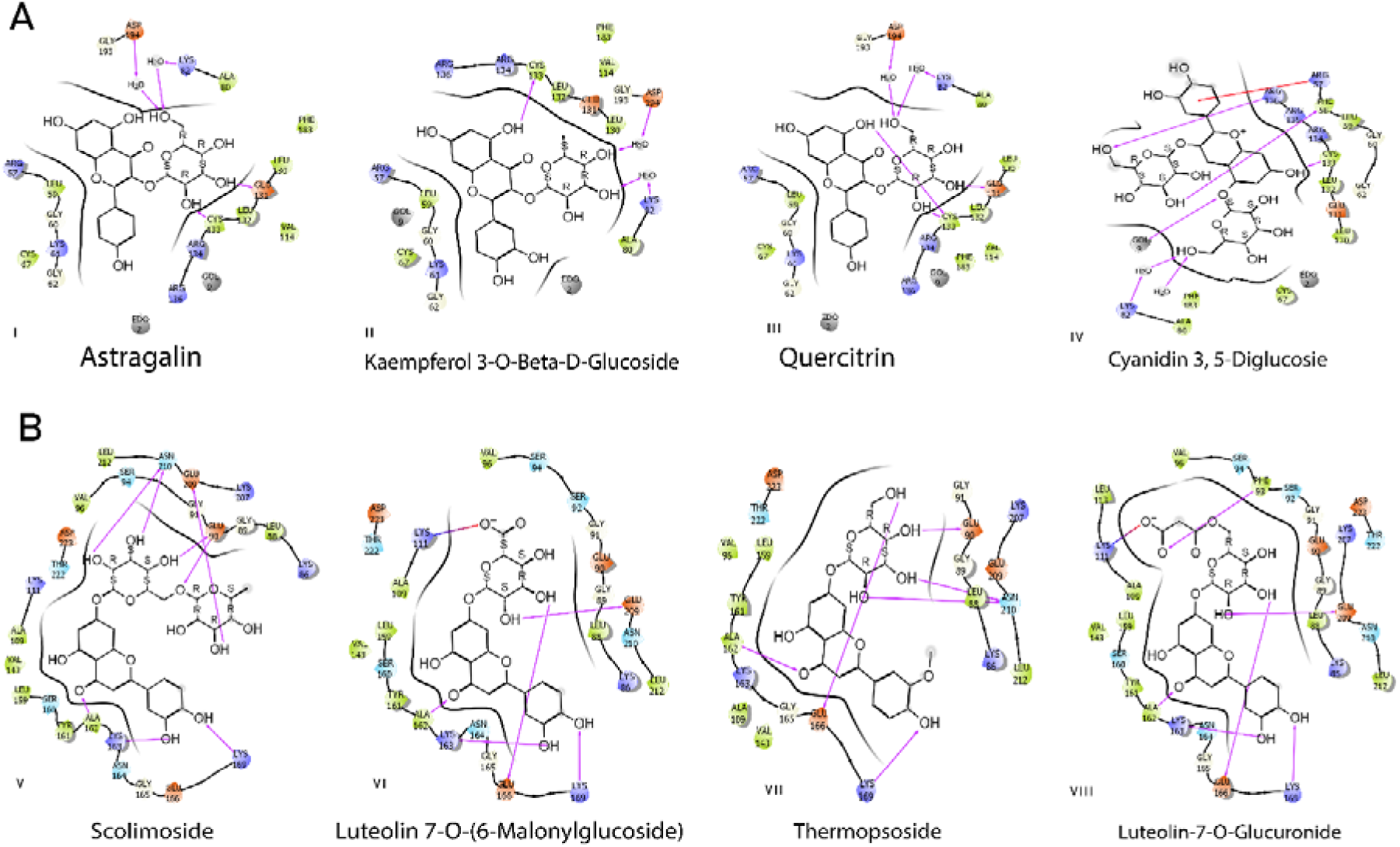
2D representation of the docked poses of selected compounds: (A) the PLK1 protein bound to astragalin (I), kaempferol 3-O-beta-D-glucoside (II), quercitrin (III), and cyanidin 3,5-digitosterol. (B) PDK1 protein bound with scolimoside (I), luteolin 7-O-(6-malonylglucoside), thermopsoside, and luteolin-7-O-glucuronide.

Scolimoside formed the most stable complex with the PDK1 protein (−10.37 kcal/mol) and showed a significant hydrogen bond interaction with the amino acid residues LYS 163, LYS 169, GLU 209, GLU 90, and ASN 210. The Ala 162 and GLU 90 residues interact with the oxygen atom of the ligand, indicating strong binding affinity^26^. By targeting PDK-1, Magozwi et al. reported that scolimoside and luteolin 7-O-(6-malonylglucoside) exhibit potential as novel therapeutic options for TNBC ^37^. Moreover, luteolin 7-O-(6-malonylglucoside), thermopsoside, luteolin-7-O-glucuronide, apigenin 7-O-glucoside, cynaroside, and apigenin-7-O-beta-D-glucopyranoside have binding energies ranging from - 9.717 to −8.839 kcal/mol against the PDK1 protein. The LYS111 residue formed a salt bridge interaction with the oxygen atoms of luteolin 7-O-(6-malonylglucoside) and luteolin-7-O-glucuronid, which implies that the interaction significantly contributed to the binding affinity and specificity to the PDK1 protein pocket. Figure 2 shows their binding post.

**Figure 2:**
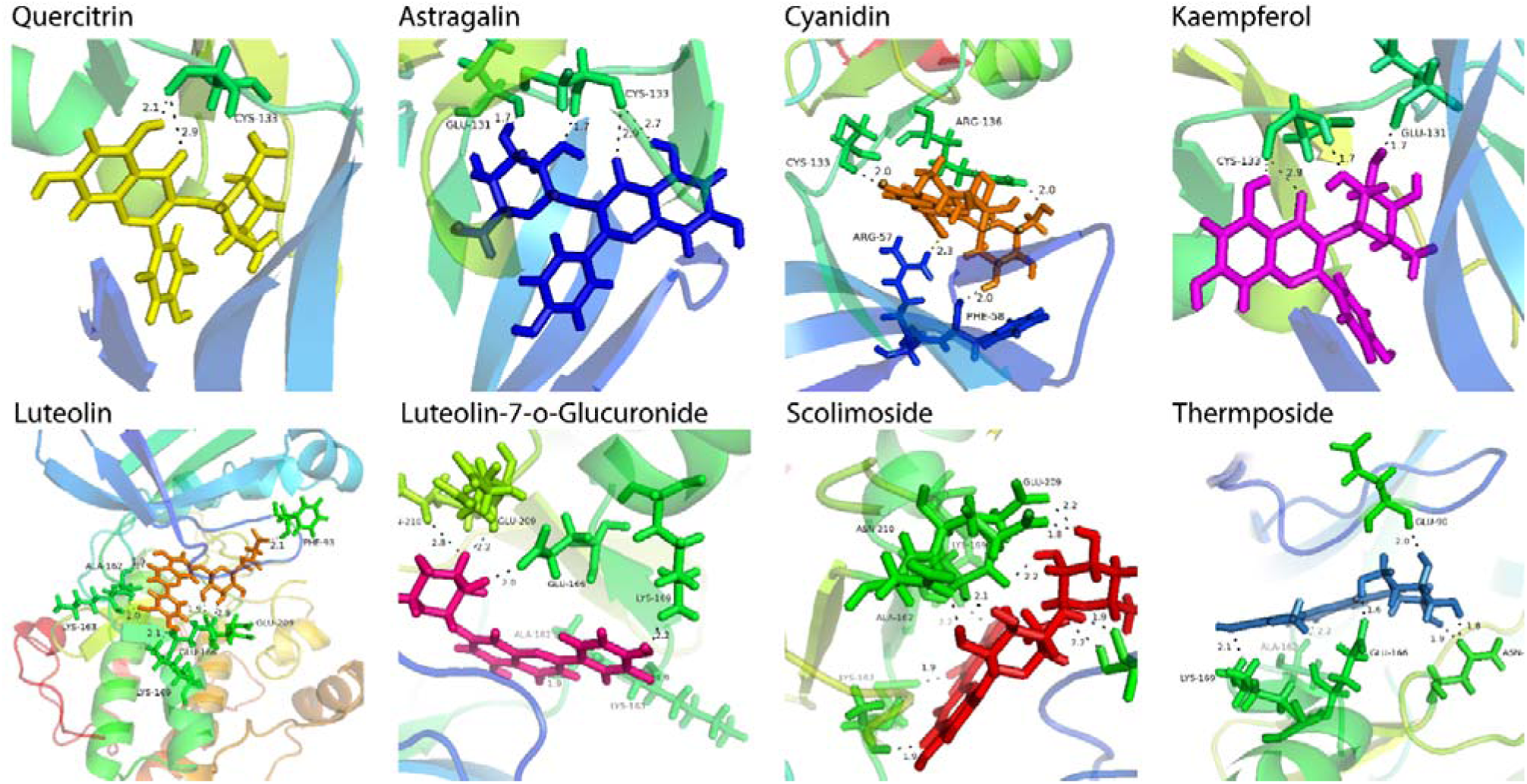
The binding poses of the all-hit compounds in the ATP active site of the PLK1 protein, namely, quercitrin (yellow), astragalin (blue), cyanidin (orange), and kaempferol (magenta), and the PDK1 protein, namely, luteolin (tv_orange), luteolin-O-glucuronide (hot pink), scolimoside (red), and thermopsoside (tv_blue).

### 3.2. ADMET analysis and free energy calculation

In addition to evaluating the binding affinity of the selected compounds for the target proteins, an ADMET analysis was also conducted to assess their drug-like properties. ADMET analysis revealed that the selected compounds possess desirable drug-like properties, including good absorption, oral bioavailability, and overall favourable pharmacokinetic profiles^38^. Lipkins’ rule of 5 was applied to assess the drug likeness of the compounds, and all of them were found to comply with this rule, indicating their potential as drug candidates (Tables 1S–2S). The toxicity properties of the ligands, including their potential to cause tumors, reproductive effects, irritation, and mutations, were evaluated using the Data Warrior software tool. None of the studied compounds displayed any toxic effects (Table 3S). This analysis indicated that the ligands have favourable toxicity profiles and can be further optimized as lead compounds for drug development. The favourable results of ADMET analysis led to further studies to gain insights into the binding free energy of these compounds and their potential as PLK1 and PDK1 inhibitors.

### 3.3. MD simulation analysis

Molecular dynamics (MD) simulations were employed to assess the stability and binding free energies of the PLK1 and PDK1 complexes with various ligands. We focused on complexes formed by astragalin, kaempferol 3-O-β-D-glucoside, and quercitrin with PLK1 and scolimoside, luteolin 7-O-(6-malonylglucoside), and thermopsoside with PDK1. Each complex was simulated for 150 ns. The root mean square deviation (RMSD) of the backbone atoms, the root mean square fluctuation (RMSF), the radius of gyration, and hydrogen bond formation were meticulously analysed for all six compounds and a reference drug in complex with both PLK1 and PDK1. Lower RMSD values indicate greater protein stability. In the PLK1 binding site, kaempferol 3-O-β-D-glucoside and astragalin exhibited average RMSD values of 0.38 nm and 0.36 nm, respectively, compared to epirubicin’s value exceeding 0.5 nm. This suggests greater binding stability for these two compounds during the simulation. Quercitrin showed moderate stability. Notably, all three PDK1 complexes (scolimoside, luteolin 7-O-(6-malonylglucoside), and thermopsoside) remained stable throughout the simulation (average RMSD values of 0.35 nm, 0.28 nm, and 0.2 nm, respectively) compared to the control. The stability of the selected compounds indicates that the active site pockets of both proteins formed quite stable interactions with the compounds. This is shown in Figure 3A&B.

**Figure 3:**
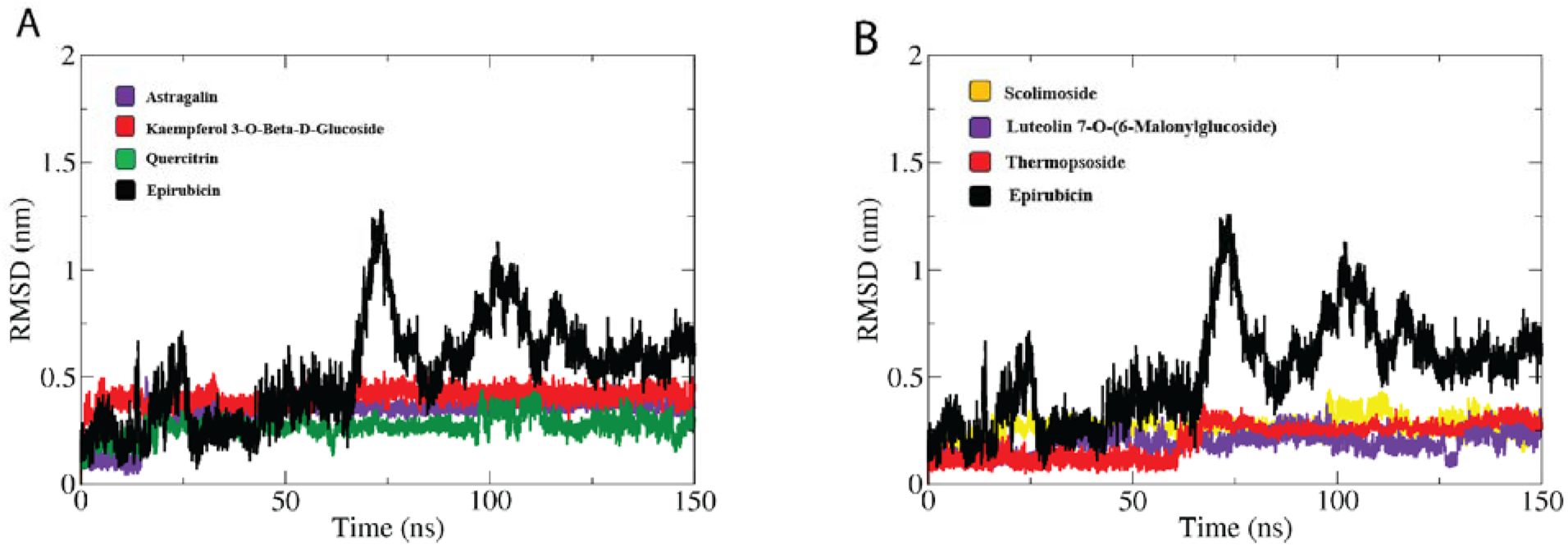
RMSD calculation for the backbone atoms in the PLK1 and PDK1 complexes. A) RMSD plot of PLK1 complexes. B) RMSD plot of PDK1 complexes.

Dynamic protein behavior was explored by analysing snapshots of protein□ligand complexes extracted at various time points (0, 25, 50, 75, 100, 125, and 150 ns) and aligning them (Figure 4). Kaempferol-3-O-β-D-glucoside and epirubicin exhibited limited stability, deviating from their initial docked positions within PLK1 and PDK1, respectively. Conversely, other compounds remained firmly bound to the binding site, indicating stable interactions. Analysis of residue dynamic behavior throughout the simulations yielded root mean square fluctuation (RMSF) values for PLK1 and PDK1, where higher values indicate greater protein flexibility. As visualized in Figure 5A-D, the C- and N-terminal regions and loop areas exhibited consistently greater overall movement across all complexes for both proteins. This suggests increased structural variability in these regions, potentially influencing protein function or interactions with ligand molecules.

**Figure 4:**
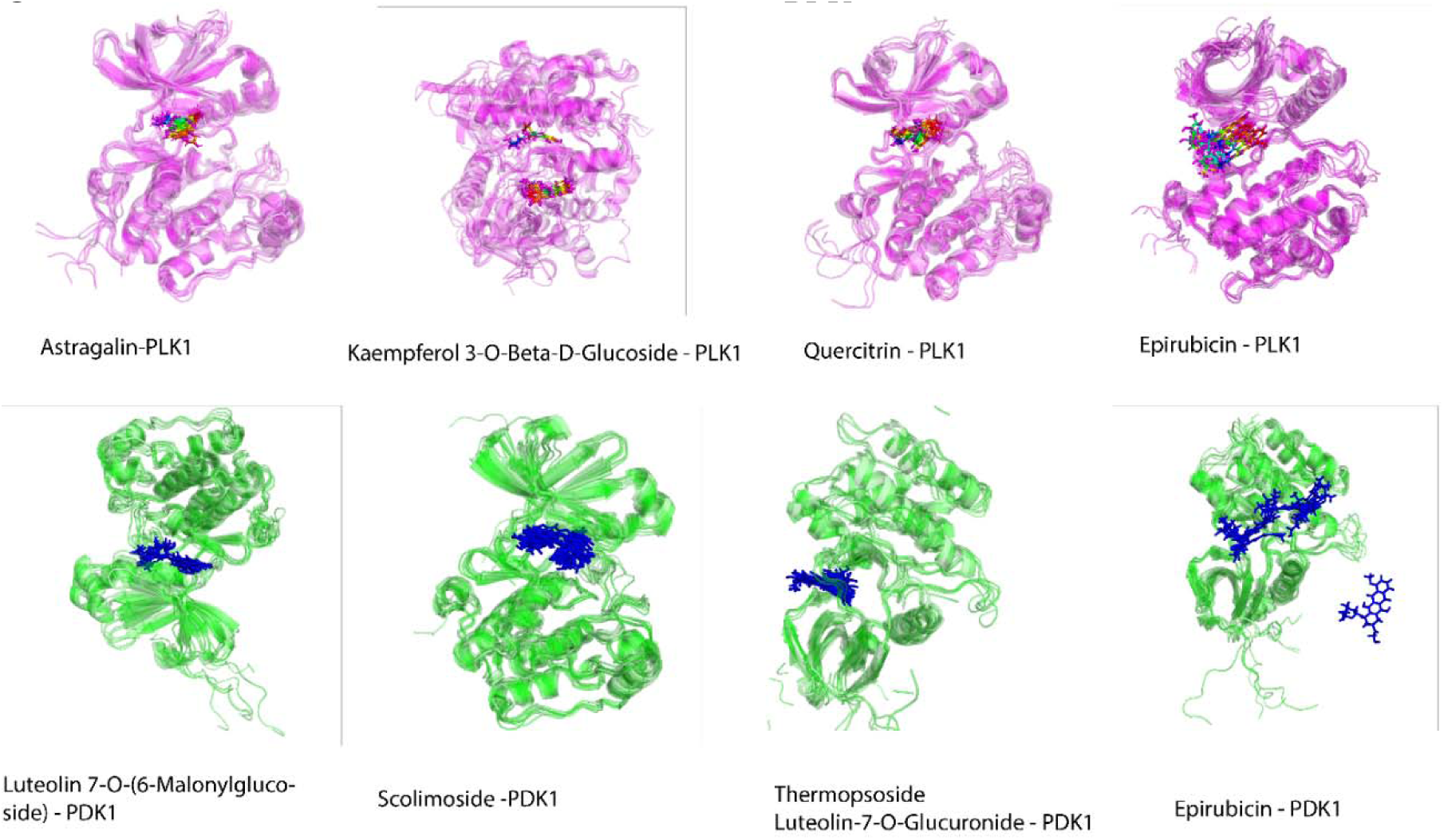
The superimposition of snapshots of PLK1 complexes and PDK1 complexes at different simulation times to reveal the dynamic and conformational changes of the selected ligands.

**Figure 5:**
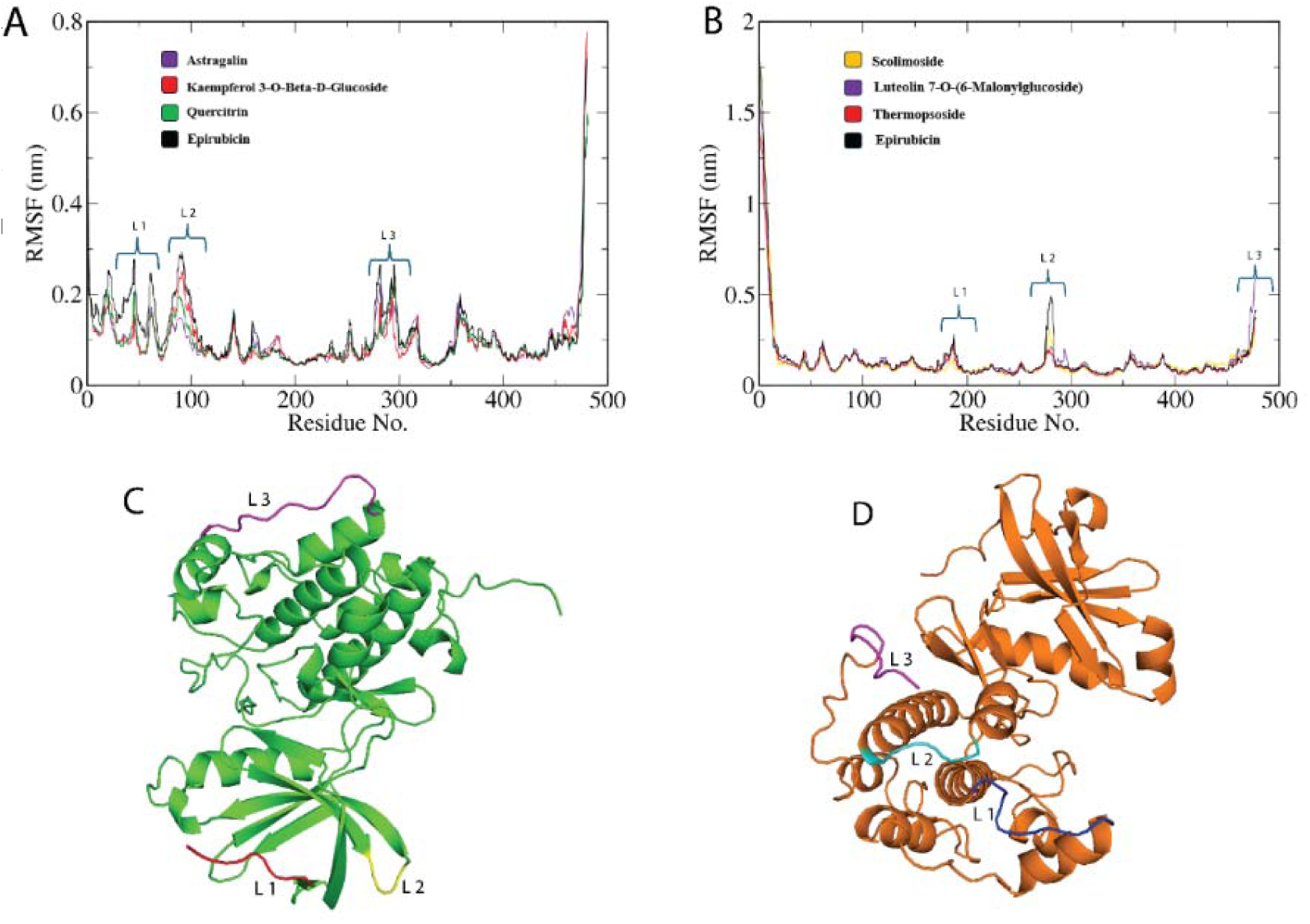
RMSF of residues in PLK1 and PDK1 during MD simulations: (A) RMSF for PLK1 complexes with astragalin, kaempferol 3-O-Beta-D-Glucoside, quercitrin, and epirubicin; (B) RMSF for PDK1 complexes with SCOLIMOSIDE, LUTEOLIN 7-o(6-MalonylGlucoside), thermopsoside, and epirubicin; (C) the structure of PLK1; (D) the structure of PDK1.

The PLK1 and PDK1 backbones exhibited distinct fluctuation patterns during the simulations. PLK1 showed structural variations in three regions (L1, L2, and L3) near its binding site, as visualized in Figures 5A and C. Interestingly, compounds such as kaempferol 3-O-β-D-glucoside and epirubicin bound to PLK1 caused higher RMSF values in L2 and L3 than astragalin and quercitrin. This suggests that astragalin and quercetin binding might restrict the movement of these regions, potentially impacting protein function or interactions with other molecules. Similarly, PDK1 displayed two prominent fluctuation regions (L1 and L2) close to its binding site (Figure 5B and D). Compounds such as luteolin 7-O-(6-malonylglucoside) and epirubicin induced significant fluctuations in L2 and L3, while leaving the loop region L1 relatively stable. This finding implies that these compounds might influence the mobility of specific regions in PDK1, potentially affecting protein interactions.

The radius of gyration (Rg) measures how the atoms are spread out around the axis of a protein. This provides information about the compactness profile of a complex in a living system ^39^. The Rg values of the control (epirubicin), astragalin, kaempferol 3-O-beta-D-glucoside, and quercitrin in complex with PLK1 were 2.05, 2.01, 2.00, and 2.03 nm, respectively. In the PDK1 complex, the control (epirubicin), scolimoside, luteolin 7-O-(6-malonylglucoside), and thermopsoside had Rg values of 2.05, 1.95, 2.06, and 2.07 nm, respectively.

In Figure 6A, the compounds astragalin, kaempferol 3-O-beta-D-glucoside, and quercitrin are shown in complex with the protein PLK1. The Rg values of these compounds were slightly lower than those of the control compound epirubicin, suggesting that they may form slightly more compact complexes with PLK1. The significantly lower Rg value of scolimoside in Figure 6B compared to epirubicin indicates a more compact structure with PDK1. This could indicate a strong and stable interaction, potentially making scolimoside an interesting compound for further study in the context of PDK1. On the other hand, luteolin 7-O-(6-malonylglucoside) and thermopsoside have slightly greater Rg values, which could mean that their complexes with PDK1 are less compact. A less compact complex may suggest weaker or more flexible binding. Hydrogen bonding analysis was further performed to evaluate the interaction strength and stability between the PLK1 complexes and PDK1 complexes. Astragalin and kaempferol 3-O-beta-D-glucoside had an average of 2-8 H-bonds with the PLK1 protein, whereas quercitrin had 2-6 H-bonds (Figure 7A). This finding implies that astragalin and kaempferol 3-O-beta-D-glucoside have stronger and potentially more stable interactions with PLK1 and might serve as potential drugs against the PLK1 protein. For the PDK1 complex, scolimoside had a large number of hydrogen bonds (3–10 H-bonds), and luteolin 7-O-(6-malonylglucoside) had a 2–8 H-bond interaction, while thermopsoside had a 1–5 H-bond interaction with the PDK1 protein (Figure 7B). A greater number and range of hydrogen bonds can be associated with more robust interactions, which might contribute to the efficacy of these compounds as inhibitors of PDK1 protein activity.

**Figure 6:**
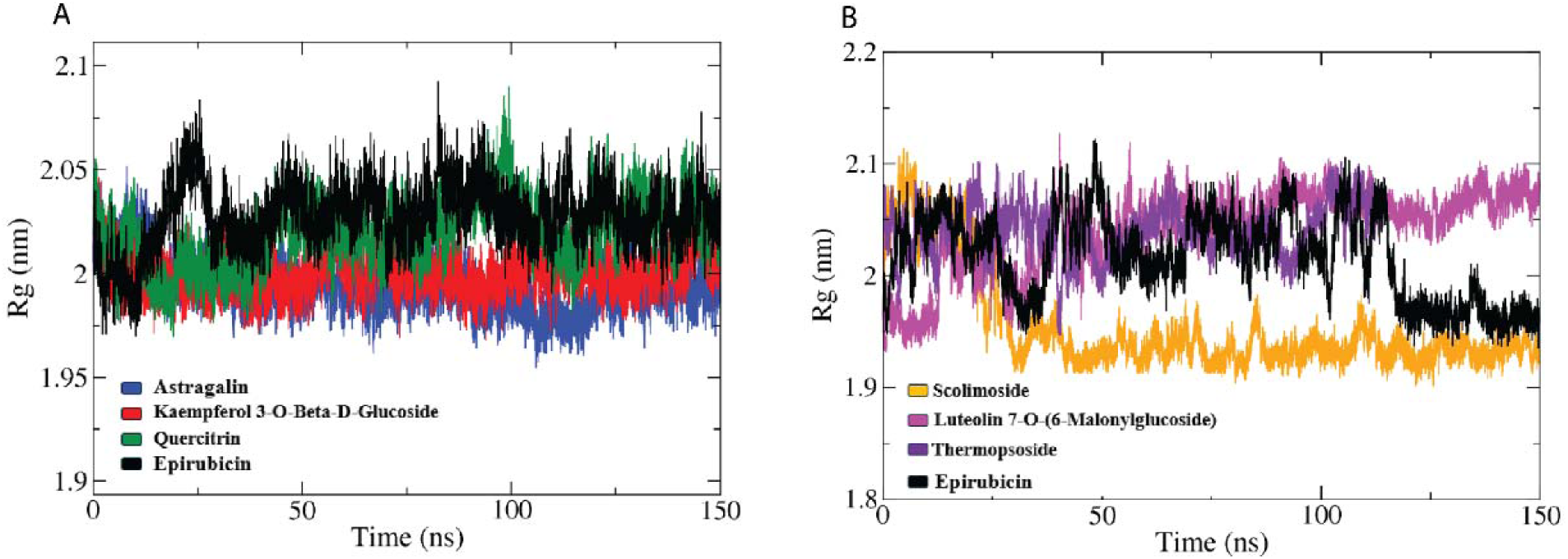
The radius of gyration (Rg) profile of the PLK1 (A) and PDK1 (B) complexes and reference compound during molecular dynamics simulations.

**Figure 7:**
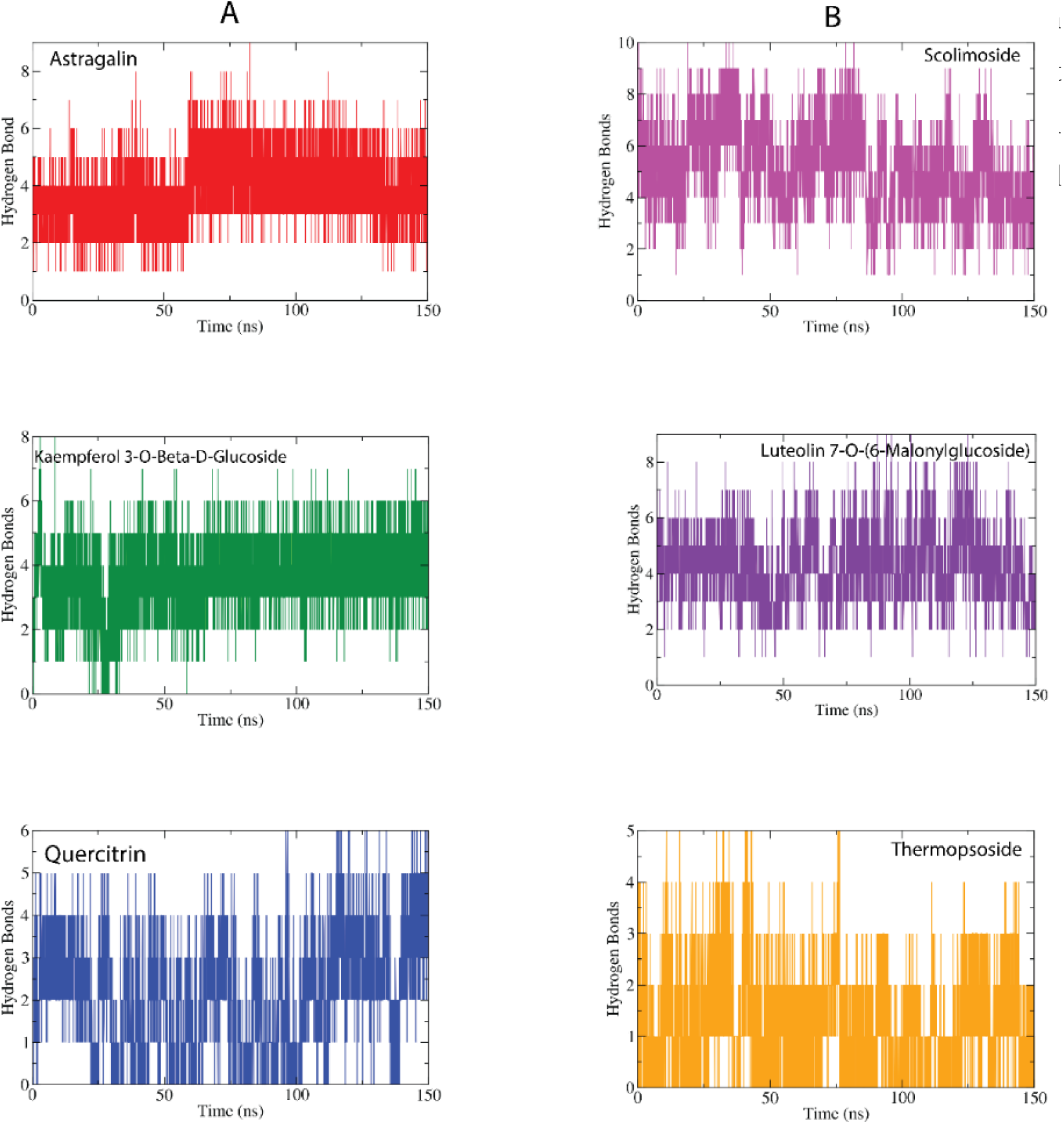
Hydrogen bond analysis of the PLK1 (A) and PDK1 (B) complexes.

### 3.4. Principal component analysis

Principal component analysis (PCA) is commonly employed to analyse coordinated movements within protein structural domains, efficiently distinguishing significant collective behaviors from ensembles derived from experimental research ^40^. In this study, PCA was applied to explore the molecular dynamics driving the selective attachment of various compounds, including astragalin, kaempferol 3-O-beta-D-glucoside, and quercitrin, to PLK1, as well as scolimoside, luteolin 7-O-(6-malonylglucoside), and thermopsoside, to PDK1. The PCA involved the creation of a covariance matrix based on the Cα atomic coordinates from the initial conformational trajectory. The proportion of variance in the first eigenvalue for kaempferol 3-O-beta-D-glucoside bound to PLK1 was 23.5%, which was greater than that of the first eigenvalue of PLK1 bound to quercitrin (19.9%) and quercitrin (19.3%), indicating greater structural variation in the kaempferol 3-O-beta-D-glucoside-PLK1 complex.

Scolimoside exhibited a greater proportion of variance in the first eigenvalue (65%) for binding to PDK1 than did the other ligands, namely, luteolin 7-O-(6-malonylglucoside) and thermopsoside, which had eigenvalues of 39.3% and 36%, respectively. The eigenvalues were calculated for 20 hyperspaces; the most significant variations were observed in the first five hyperspaces before reaching a constant. The overall flexibility of the first three PCs was calculated, and for PKL1, quercitrin exhibited the highest variability (46.65%) as shown in figure 8, followed by astragalin and kaempferol 3-O-beta-D-glucoside, with variabilities of 43.2% and 44.07%, respectively, (as depicted in figure 2S). In the case of PDK1, scolimoside (figure 9) bound to PDK-1 had greater variability (84.02%) than luteolin 7-O-(6-malonylglucoside) (72.94%) and thermopsoside (74.03%), which are respectively shown in figure 3S. Grouping the data points in the principal component space identified distinct conformations within each cluster. Notably, the blue regions exhibited the most pronounced structural shifts, while the white regions showed moderate changes. In contrast, the red regions displayed minimal movement or flexibility.

**Figure 8:**
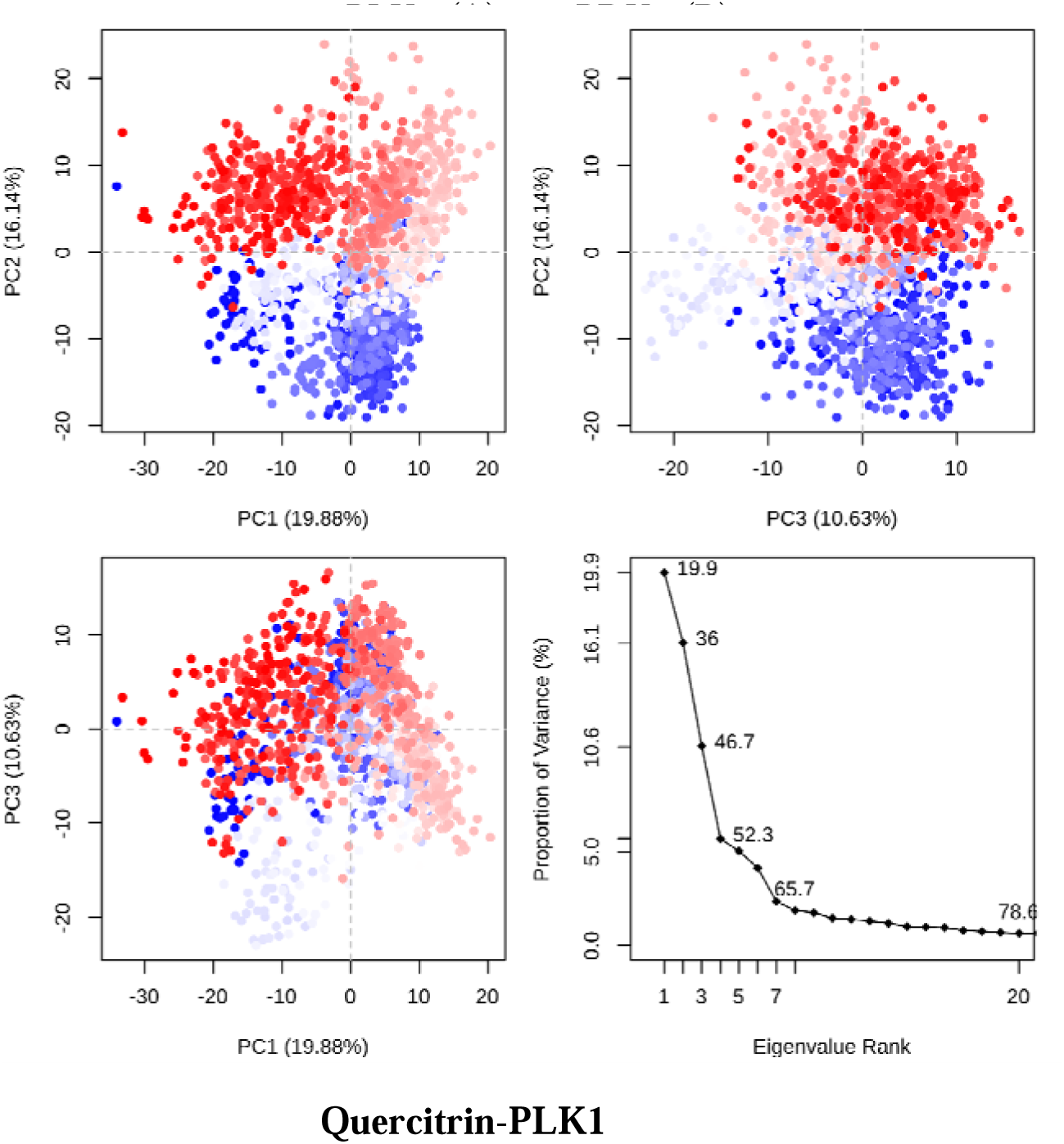
Graphical representation of the percentage of variance in the PLK1 complex. The fluctuation regions are shown in the three PCs: PC1, PC2, and PC3. The overall fluctuations for the Quercitrin-PLK1 complex (46.65%)

**Figure 9:**
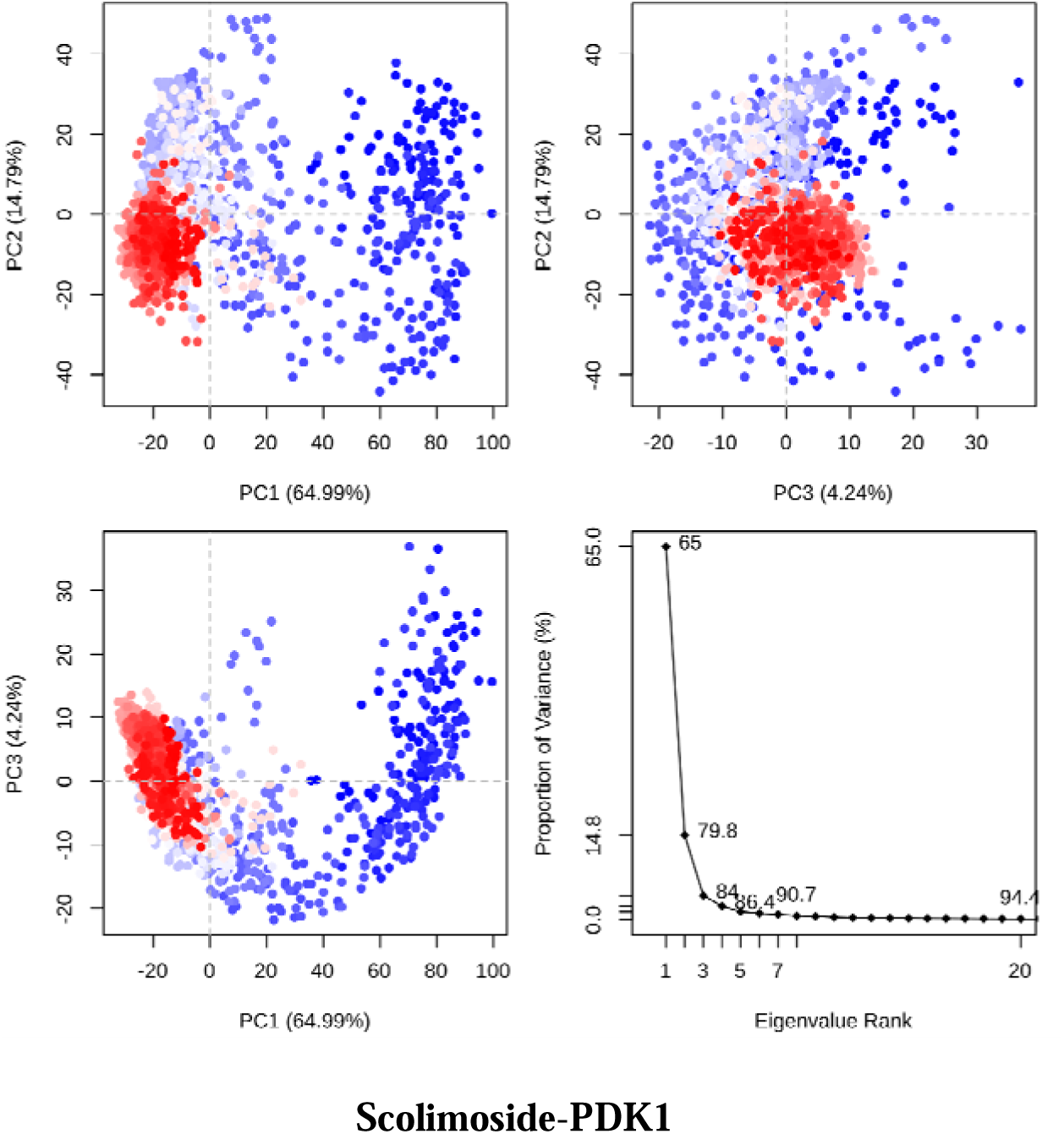
Graphical representation of the percentage of variance in the PDK1 complex. The fluctuation regions are shown in the three PCs: PC1, PC2, and PC3. The overall fluctuations for the solimoside-PDK1 complex (84.02%, 44.07%),

### 3.5. Dynamic Cross-Correlation Network of PLK1 and PDK1 proteins

The comparative structural dynamics between the PLK1 and PDK1 complexes were analysed by calculating the cross-correlation matrices for the Cα atomic coordinates, aiming to identify variances in their structural behaviours ^41^. Cyan and purple hues were utilized to represent positively and negatively correlated movements, respectively. The matrix’s diagonal section highlights the self-correlated movements of each domain, whereas interactions between different domains are illustrated in the nondiagonal areas. The straight line running from the lower left corner to the upper right, which is consistently colored, denotes the self-correlation of individual residues, invariably scoring +1.0, indicating a perfect positive correlation ^42^. The binding of compounds such as astragalin, kaempferol 3-O-beta-D-glucoside, quercitrin, scolimoside, luteolin 7-O-(6-malonylglucoside), and thermopsoside significantly impacted the structural dynamics of PLK1 and PDK1.

For the PLK1 complex (Figure 10A, 10B, 10C), anti-correlations were observed in the R2 and R3 regions, while positive correlations were observed in the diagonal and R1 regions. In the case of the PDK1 complex (Figure 10D, 10E, 10F), a positive correlation motion was observed in regions R1, R2 and the diagonal, while an anti-correlation motion was prominent in regions R3 and R4. These findings suggest that the tested compounds can induce distinct structural changes in PLK1 and PDK1, potentially affecting their function and signalling pathways.

**Figure 10:**
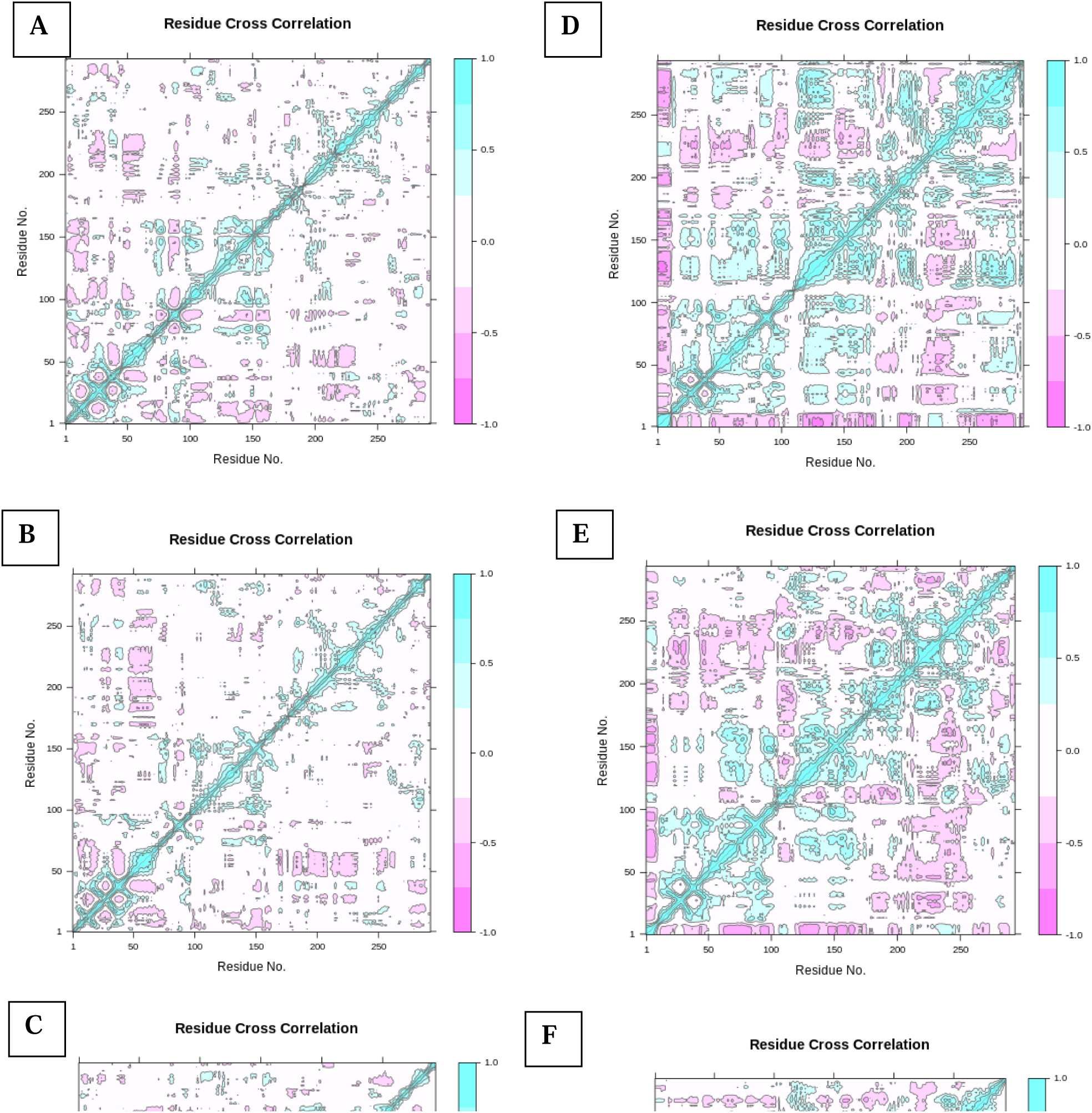
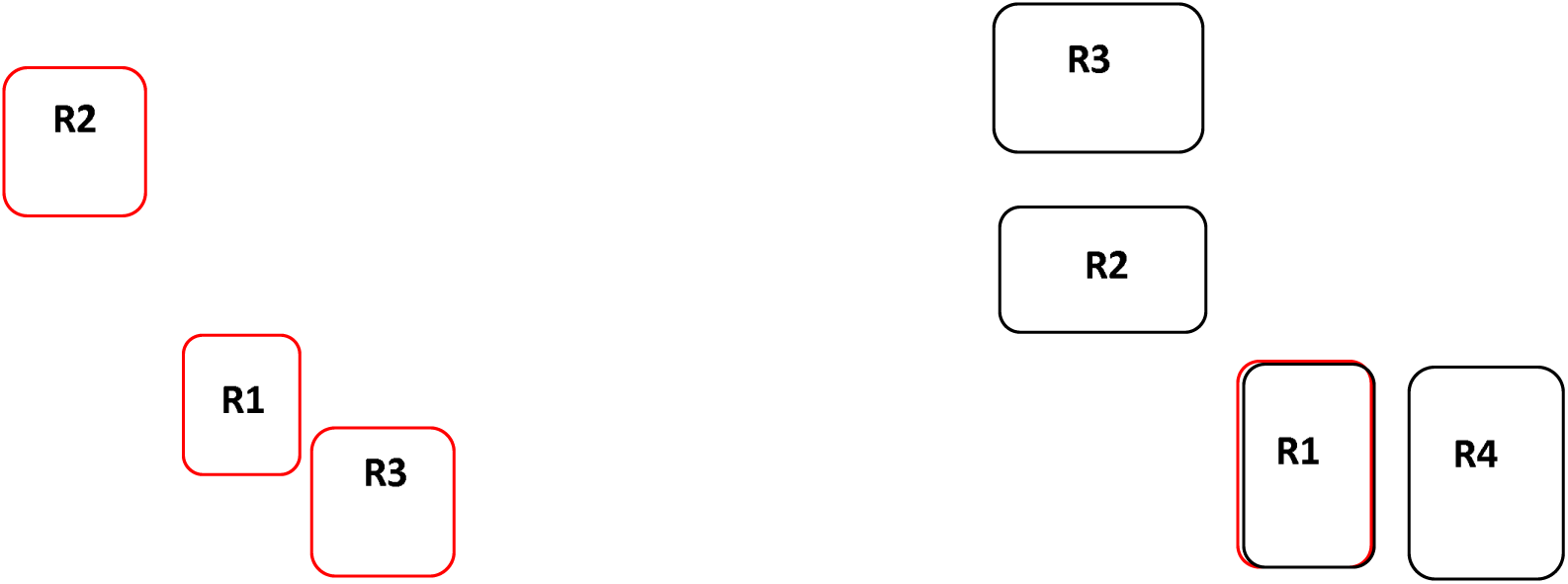
Dynamic cross-correlation map for the Cα atom pairs within the PLK1 complex: astragalin (A), kaempferol 3-O-beta-D-glucoside (B), and quercitrin (C); and the PDK1 complex: scolimoside (D), luteolin 7-O-(6-malonylglucoside) (E), and thermopsoside (F).

### 3.6. Binding free energy calculation (MM-GBSA) studies

The gmx_MMPBSA method ^43^ was used to test how well the chosen hit compounds bound to the PLK1 and PDK1 receptors. The MMGBSA method was used due to its computational efficiency, which allows for faster free energy calculations without significantly compromising accuracy. Given the size and complexity of the protein-ligand systems involved, MMGBSA was selected as a more practical approach for this study, providing a balance between speed and reliable predictions of binding affinities. A more negative Δ_TOTAL_ value signifies a stronger binding affinity ^44^. For both PLK1 and PDK1, the calculations were performed by summing the free energies of the protein-ligand complex and removing the free energies of the separate proteins and ligands. This total binding free energy incorporates contributions from van der Waals interactions, electrostatic interactions (EEL), and the generalized born (GB) solvation term (EGB) ^45^.

Among the investigated compounds, astragalin, kaempferol 3-O-β-D-glucoside, and quercetin displayed significantly more favourable binding affinities toward PLK1 complexes than did the reference drug epirubicin. This was evidenced by their more negative ΔVDWAALS values (−48.46 ± 0.06, −49.54 ± 0.07, and −42.28 ± 0.07 kcal/mol, respectively) compared to those of epirubicin (−38.67 ± 0.10 kcal/mol). Notably, astragalin exhibited the strongest van der Waals interactions (ΔVDWAALS = −48.46 ± 0.06 kcal/mol) and the most negative ΔEEL value (−52.55 ± 0.15 kcal/mol), indicating that the most favourable electrostatic interactions occurred among the ligands. Kaempferol 3-O-β-D-glucoside had the highest surface energy binding affinity of −6.81 ±0.00 kcal/mol, while epirubicin had the lowest. The free energy in the gas phase (ΔGGAS) combines van der Waals and electrostatic energies. This is highly favourable for all the selected ligands, but epirubicin is unfavourable due to its positive value. Interestingly, epirubicin had the most positive ΔEGB and ΔGSOLV values, suggesting unfavourable solvation that counteracts other favourable interactions. For the total binding free energy, astragalin exhibited the most favourable binding affinity towards the PLK1 protein due to a combination of factors, as evidenced by the data in Table 3A.

**Table 3:** The binding free energy with gmxMMPSA for the calculation of the MMGBSA of hit compounds of Daucus carota against 3FC2 (A) and 4AW1 (B).

| **A. PLK1 Protein (3FC2)** |  |  |  |  |
| --- | --- | --- | --- | --- |
|  | Astragalín | Kaempferol 3-O-Beta-D-Glucoside | Quercitrín | Epirubicín |
|  | Average (Kcal/mol) | Average (Kcal/mol) | Average (Kcal/mol) | Average (Kcal/mol) |
| $\Delta_{\text{VDWAALS}}$ | $-48.46 \pm 0.06$ | $-49.54 \pm 0.07$ | $-42.28 \pm 0.07$ | $-38.67 \pm 0.10$ |
| $\Delta_{\text{EEL}}$ | $-52.55 \pm 0.15$ | $-49.27 \pm 0.16$ | $-30.74 \pm 0.33$ | $185.30 \pm 0.25$ |
| $\Delta_{\text{EGB}}$ | $55.94 \pm 0.11$ | $54.70 \pm 0.11$ | $42.68 \pm 0.20$ | $-158.62 \pm 0.23$ |
| $\Delta_{\text{ESURF}}$ | $-6.72 \pm 0.00$ | $-6.81 \pm 0.00$ | $-6.24 \pm 0.01$ | $-4.87 \pm 0.01$ |
| $\Delta_{\text{GGAS}}$ | $-100.81 \pm 0.15$ | $-98.81 \pm 0.15$ | $-73.03 \pm 0.30$ | $147.04 \pm 0.27$ |
| $\Delta_{\text{GSOLV}}$ | $49.22 \pm 0.11$ | $47.89 \pm 0.11$ | $36.43 \pm 0.20$ | $-163.49 \pm 0.23$ |
| $\Delta_{\text{TOTAL}}$ | $-51.58 \pm 0.07$ | $-50.91 \pm 0.09$ | $-36.59 \pm 0.15$ | $-16.45 \pm 0.09$ |
| **B. PDK1 Protein (4AW1)** |  |  |  |  |
|  | Scolimoside | Luteolín 7-O-(6-Malonylglucoside) | Thermopsoside | Epirubicín |
|  | Average | Average | Average | Average |
| $\Delta_{\text{VDWAALS}}$ | $-51.51 \pm 0.08$ | $-43.96 \pm 0.07$ | $-44.94 \pm 0.07$ | $-14.19 \pm 0.10$ |
| $\Delta_{\text{EEL}}$ | $-69.26 \pm 0.28$ | $-129.51 \pm 0.60$ | $-51.00 \pm 0.25$ | $-81.35 \pm 0.75$ |
| $\Delta_{\text{EGB}}$ | $75.63 \pm 0.17$ | $146.96 \pm 0.39$ | $59.50 \pm 0.16$ | $77.82 \pm 0.66$ |
| $\Delta_{\text{ESURF}}$ | $-7.95 \pm 0.01$ | $-7.04 \pm 0.01$ | $-6.54 \pm 0.01$ | $-2.46 \pm 0.01$ |
| $\Delta_{\text{GGAS}}$ | $-120.77 \pm 0.26$ | $-173.47 \pm 0.60$ | $-95.95 \pm 0.23$ | $-95.16 \pm 0.75$ |
| $\Delta_{\text{GSOLV}}$ | $67.68 \pm 0.17$ | $139.91 \pm 0.51$ | $52.96 \pm 0.16$ | $75.37 \pm 0.65$ |
| $\Delta_{\text{TOTAL}}$ | $-53.09 \pm 0.15$ | $-33.55 \pm 0.15$ | $-42.99 \pm 0.11$ | $-13.95 \pm 0.12$ |

Analysis of protein□ligand interactions for PDK1 revealed that scolimoside exhibited the most favourable van der Waals interactions (ΔVDWAALS = −51.51 ± 0.08 kcal/mol) and greatest surface energy (ΔESURF = −7.95 ± 0.01 kcal/mol). In contrast, luteolin 7-O-(6-malonylglucoside) displayed the most electrostatic interactions (ΔEEL = −129.51 ± 0.60 kcal/mol). These findings collectively demonstrate favourable binding of all flavonoids to both the PLK1 and PDK1 proteins, with varying degrees of binding free energy (Δ_TOTAL_). Notably, astragalin and scolimoside exhibited the most favourable Δ_TOTAL_ values for PLK1 and PDK1, respectively. Interestingly, epirubicin displayed a distinct profile characterized by highly unfavourable electrostatic interactions, compensated by favourable solvation energies. Consequently, epirubicin exhibited lower negative Δ_TOTAL_ values, suggesting a weaker binding affinity for both proteins than for flavonoids. The contributions of each energy to the total binding energy are shown in Table 3B.

To further ascertain the impact of crucial residue interactions with hit ligands in terms of the total binding free energy contribution and favourable binding affinities, a per-residue energy decomposition analysis ^46^ was conducted. This analysis breaks down the total binding free energy into contributions from individual residues, allowing the identification of key interaction points and residues critical for ligand binding affinity, as depicted in Table 4A&B ^46^. The decomposition analysis showed that the hit ligands interacted with both PLK1 and PDK1. The PLK1 residues PHE183 (4.22 ± 0.04 kcal/mol), CYS67 (2.26 ± 0.03 kcal/mol), and LEU59 (2.26 ± 0.04 kcal/mol) had favourable contributions to the total binding free energy for all the PLK1 protein□ligand interaction complexes. The side chain of the PHE183 residue significantly contributed to the binding affinity of kaempferol 3-O-Beat-D-glucoside for the PLK1 protein. In the complexes with astragalin and kaempferol 3-O-beta-D-glucoside, the CYS133 residue (−1.76 ± 0.04) also contributed favourably. However, the LEU132 residue had an unfavourable interaction with kaempferol 3-O-beta-D-glucoside due to this interaction having positive binding free energy values. Residue ASP194 showed a favourable interaction with quercetin (−3.48 ± 0.21 kcal/mol).

**Table 4:**
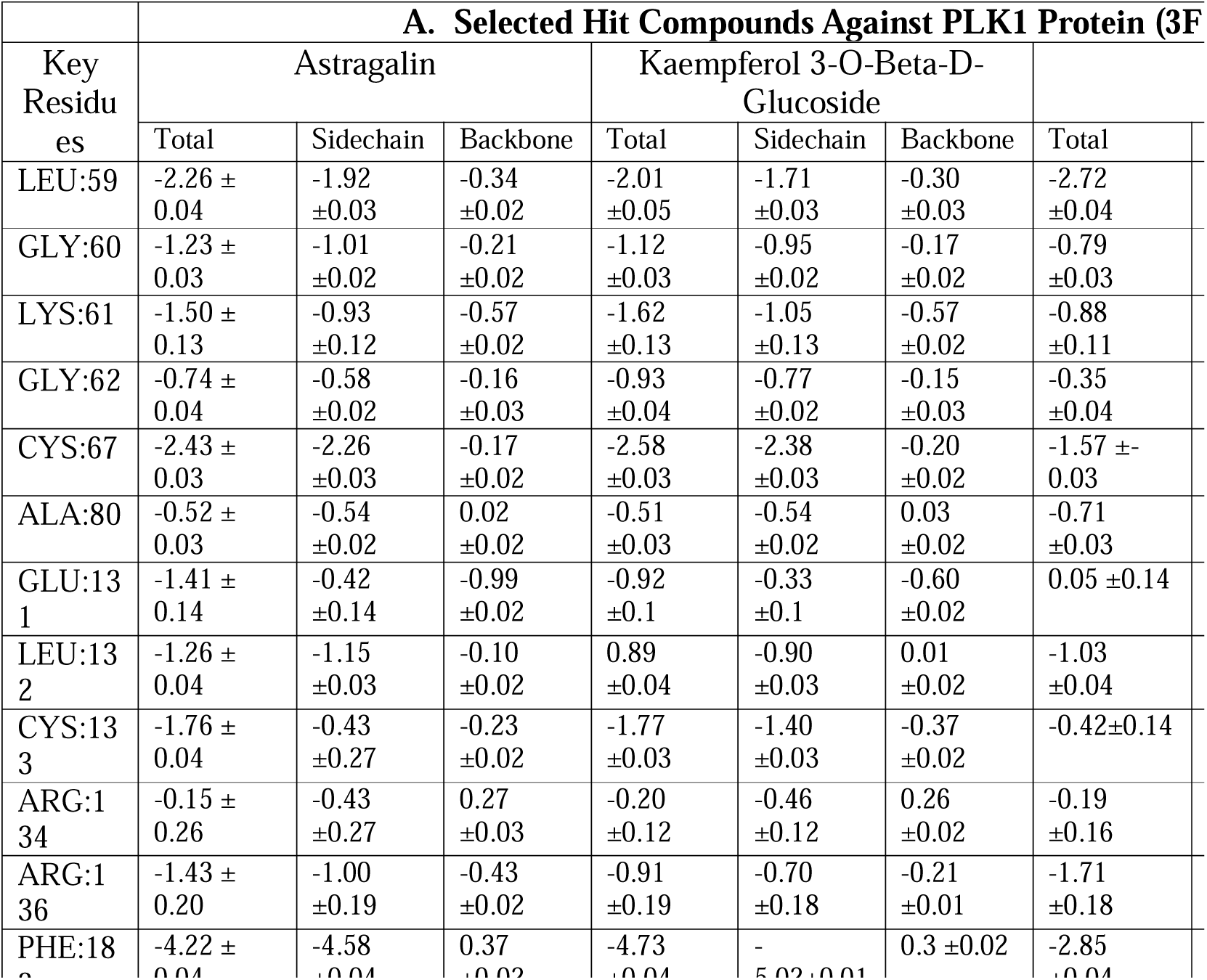
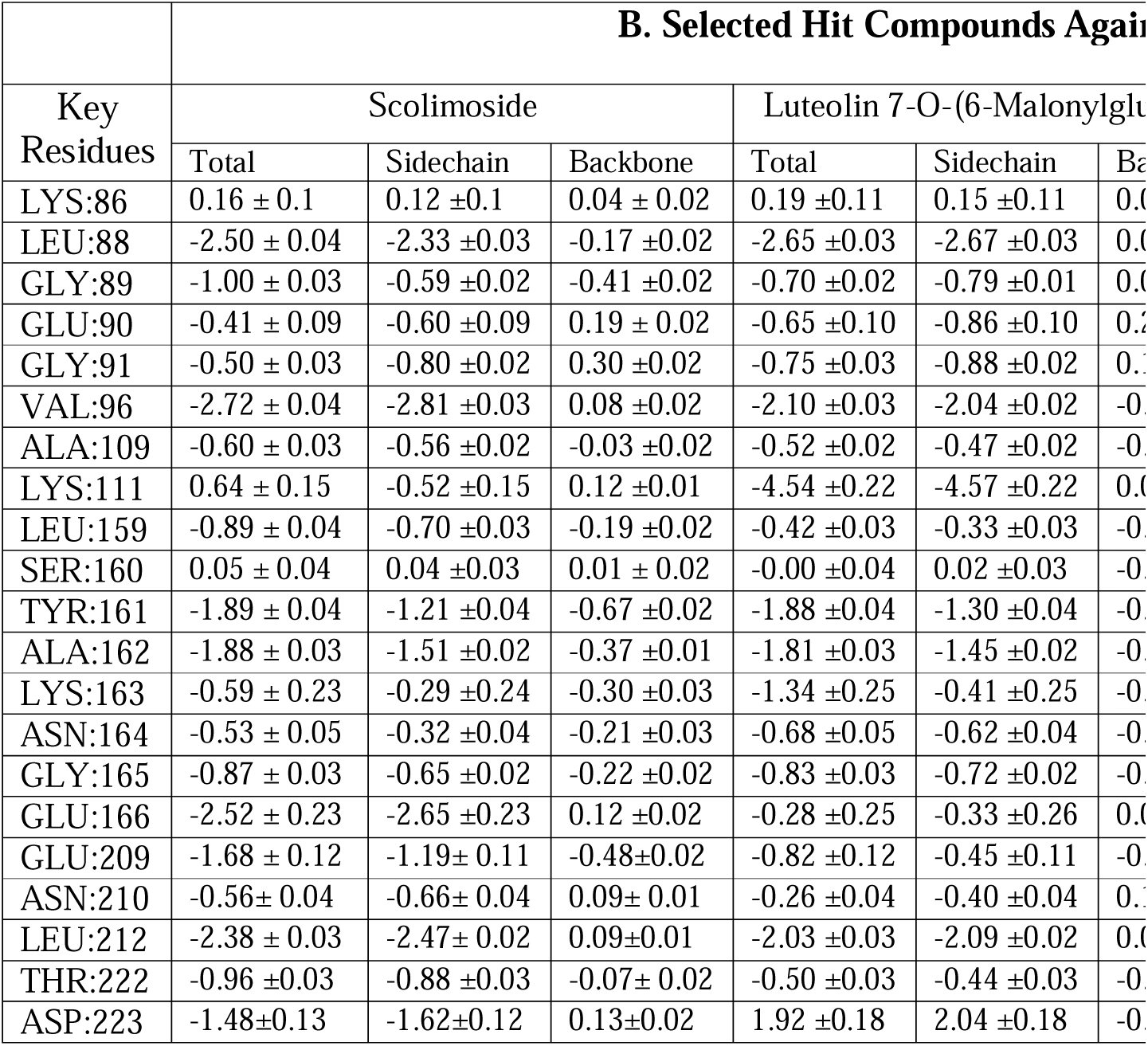
Analysis of Energy Decomposition: This involves the breakdown of ΔG_ligand– residue values into the individual contributions from the side chains and the backbone of crucial residues within PKL1 (A) and PDK1 (B).

On the other hand, the PDK1 residues LEU88 and VAL96 were strongly negatively related to the total binding free energy across all three compounds, indicating favourable interactions. Specifically, LEU88 shows a consistent and significant negative energy contribution, underscoring its importance in the binding site architecture and its role in stabilizing interactions with ligands. LYS111 exhibits a particularly interesting interaction profile with luteolin 7-O-(6-malonylglucoside), showing a dramatic shift to a highly favourable interaction (−4.54 ± 0.22 kcal/mol) compared to its interaction with the other two compounds (Figure 12). This suggests that LYS111 may have a unique interaction mechanism with Luteolin 7-O-(6-malonylglucoside) that significantly contributes to binding affinity, potentially through specific hydrogen bonding or electrostatic interactions. GLU166 and LEU212 both showed favourable contributions across different compounds, although the magnitude of their interactions varied. GLU166, for example, shows a markedly less favourable interaction with luteolin 7-O-(6-malonylglucoside) than with scolimoside and thermopsoside. This variability underscores the nuanced role these residues may play in ligand specificity and affinity^47^. However, the interaction of the ASP223 residue with luteolin 7-O-(6-malonylglucoside) has an unusual positive effect, suggesting an unfavourable interaction. This could be due to destabilizing effects in the ligand-residue interaction ^48^.

### 3.7. Alanine scanning mutagenesis of selected complexes

The PLK1 and PDK1 complexes were further analysed to provide insight into the key roles of the residues involved in the total binding free energy of protein□ligand complexes via computational alanine scanning mutagenesis ^49^. This method involves systematically mutating each residue of interest to alanine (except when alanine is the original residue, in which case it is mutated to glycine) to assess the impact of each residue’s side chain on the binding affinity of the ligand to the protein^50^. This computational technique provides a quantitative measure of the change in binding free energy (ΔΔGbind) resulting from the mutation, offering insights into the contribution of individual residues to the overall stability and specificity of the protein□ligand interaction ^51 49^. The contributions of the CYS133, ASP194, and GLU131 residues of PLK1 and the Ala162 (mutated to glycine), LYS169, GLU166, ASN210, and LYS163 residues of PDK1 were estimated one by one. The change in the binding free energy of the complexes can be observed in Tables 5A and 5B.

**Table 5:** Alanine Scanning: Calculation of the binding free energy (Kcal/mol) of Delta mutant complexes with key residues. * WΔ_TOTAL_, binding free energy of the wild-type complex; MΔ_TOTAL_, binding free energy of the mutant complex; ΔΔ_TOTAL_, difference in mutant and wild-type energy values.

A. PLK1 Complexes with Key Residues
| Complexes | Key Residues | $\Delta_{VDWAALS}$ | $\Delta_{EEL}$ | $\Delta_{EGB}$ | $\Delta_{ESURF}$ | $\Delta_{GGAS}$ | $\Delta_{GSOLV}$ | $W\Delta_{TOTAL}$ | $M\Delta_{TOTAL}$ | $\Delta\Delta_{TOTAL}$ |
| --- | --- | --- | --- | --- | --- | --- | --- | --- | --- | --- |
| PLK1-Astragalin | C133A | -48.31±3.75 | -17.71±4.04 | 22.34± 3.16 | -6.71±0.32 | -66.02±5.58 | 15.63± 2.96 | -51.58±0.07 | -50.39±3.67 | -1.19±0.22 |
|  | D194A | -50.07±3.44 | -4.99±2.18 | 9.05±1.61 | -6.43±0.32 | -55.06±4.21 | 2.62±1.43 | -51.58±0.07 | -52.44±3.50 | -0.86±1.32 |
|  | E131A | -48.20±3.76 | -10.76±2.39 | 13.09±1.75 | -6.71±0.32 | -58.96±4.50 | 6.38 ± 1.55 | -51.58±0.07 | -52.58±3.71 | -1.00±0.06 |
| PLK1-Kaempferol 3-O-Beta-D-Glucoside | C133A | -49.65 ±4.17 | -16.49 ±3.16 | 21.86 ±2.74 | -6.81 ±0.30 | -66.13 ±4.78 | 15.05 ±2.59 | -50.91 ±0.09 | -51.08 ±3.47 | -0.17±0.18 |
|  | D194A | -51.29±3.88 | -4.96±1.58 | 9.36±1.19 | -6.58±0.29 | -56.25±3.78 | 2.79±1.10 | -50.91 ±0.09 | -53.47±3.55 | -2.56±1.25 |
|  | E131A | -49.50±4.17 | -9.91±1.85 | -9.91±1.85 | -6.81±0.30 | 12.67±1.46 | 5.87±1.33 | -50.91 ±0.09 | -53.55±3.73 | -2.64±0.02 |
| PLK1-Quercitrin | C133A | -42.29±4.41 | -10.51±6.84 | 17.02±4.75 | -6.23±0.37 | -52.81 ±5.25 | 10.79±4.76 | -36.59±0.15 | -42.02±3.25 | -5.43±0.08 |
|  | D194A | -44.59±3.12 | 0.49±1.53 | 5.37±1.21 | -5.95±0.38 | -44.10±2.71 | -0.59±1.29 | -36.59±0.15 | -44.69±2.98 | -8.1±1.59 |
|  | E131A | -38.30±353.83 | -6.71± 4.00 | 10.30± 2.68 | -6.24±0.37 | -45.82±366.75 | 4.06 ±2.69 | -36.59±0.15 | -41.76 ±366.60 | -5.17±0.02 |

B. PDK1 Complexes with Key Residues
| Complexes | Key Residues | $\Delta_{VDWAALS}$ | $\Delta_{EEL}$ | $\Delta_{EGB}$ | $\Delta_{ESURF}$ | $\Delta_{GGAS}$ | $\Delta_{GSOLV}$ | $W\Delta_{TOTAL}$ | $M\Delta_{TOTAL}$ | $\Delta\Delta_{TOTAL}$ |
| --- | --- | --- | --- | --- | --- | --- | --- | --- | --- | --- |
| PDK1-Scolimoside | A162G | -51.20±4.53 | -69.06±16.15 | 88.05±11.43 | -7.94 ±0.56 | - 120.26±16.03 | 80.12±11.21 | -53.09 ±0.15 | -40.15±6.97 | 12.49±0.38 |
|  | K169A | -50.54±4.38 | -14.56±3.01 | 16.75±1.92 | -7.78 ±0.52 | -65.11 ±4.84 | 8.971±.77 | -53.09 ±0.15 | -56.14 ±4.50 | -3.05±0.54 |
|  | E166A | -53.03±4.25 | -7.30±3.53 | 11.89±2.52 | -7.46±0.52 | -60.33±5.01 | 4.43±2.37 | -53.09 ±0.15 | -55.90±4.37 | -2.81±1.54 |
|  | K163A | -51.45 ±4.54 | -13.95±3.30 | 16.76± 2.23 | -7.95±0.56 | -65.39± 5.18 | 8.81± 2.05 | -53.09 ±0.15 | -56.58±4.65 | -3.49±0.03 |
|  | N210A | -51.17±4.46 | -22.81±5.24 | 28.24± 3.56 | -7.92±0.56 | -73.98±6.18 | 20.33±3.38 | -53.09 ±0.15 | -53.65±4.58 | -0.56±0.28 |
| PDK1-Luteolin 7-O-(6-Malonylglucoside) | A162G | -43.34±4.55 | - 123.51±37.37 | 138.64±32.77 | -6.99±0.58 | - 166.85±35.93 | 131.65±32.77 | -33.55±0.15 | -35.20±5.95 | -1.65±0.32 |
|  | K169A | -48.09±4.47 | -22.56±7.62 | 24.18±6.46 | -6.93±0.58 | -65.65±7.24 | 17.25±6.48 | -33.55±0.15 | -48.40±4.21 | -14.85±0.33 |
|  | E166A | -43.26±4.34 | -28.24±6.88 | 30.44±5.88 | -6.68±0.53 | -71.50±6.62 | 23.76±5.92 | -33.55±0.15 | -47.74±3.95 | -14.19±0.97 |
|  | K163A | -43.53±4.56 | -23.64±7.47 | 25.51±6.25 | -7.00±0.58 | -67.17±7.18 | 18.51±6.27 | -33.55±0.15 | -48.66±4.24 | -15.11±0.05 |
|  | N210A | -43.48±4.49 | -43.40±12.37 | 47.41±10.57 | -6.99±0.58 | -86.88±11.49 | 40.42±10.58 | -33.55±0.15 | -46.46±4.00 | -12.91±0.22 |
| PDK1-Thermopsoside | A162G | -44.64±4.14 | -50.63±14.60 | 68.36±11.47 | -6.53±0.44 | -95.27±13.85 | 61.83±11.29 | -42.99 ±0.11 | -33.44±4.90 | 9.55±0.31 |
|  | K169A | -44.17±4.09 | -11.02±2.95 | 13.21±2.11 | -6.44±0.43 | -55.19±4.10 | 6.77±2.01 | -42.99 ±0.11 | -48.43±3.71 | -5.44±0.50 |
|  | E166A | -44.30±3.86 | -6.95±3.02 | 10.13±2.23 | -6.18±0.43 | -51.25±3.97 | 3.95±2.13 | -42.99 ±0.11 | -47.30±3.54 | -4.31±1.17 |
|  | K163A | -44.87±4.15 | -10.19±2.94 | 12.94±2.22 | -6.54±0.44 | -55.06±4.27 | 6.40±0.00 | -42.99 ±0.11 | -48.66±3.75 | -5.67±0.03 |
|  | N210A | -44.71±4.09 | -16.94±4.87 | 22.14±3.68 | -6.51±0.43 | -61.65±5.14 | 15.62±3.53 | -42.99 ±0.11 | -46.03±3.40 | -3.04±0.31 |

In the PLK1-quercitrin complex, the mutation of ASP194 to alanine significantly altered the binding free energy of the complex, reflecting a substantial impact on the interaction dynamics between the quercitrin and PLK1 proteins. This mutation led to a considerable change in the overall binding energy, as evidenced by the data presented in Table 5A, which shows a ΔΔ_TOTAL_ of −8.10 ± 1.59 kcal/mol. This indicates that ASP194 contributes significantly to the interactions within the complex, which are crucial for the structural integrity and tight binding of quercitrin to PLK1. Similarly, the contribution of GLU131 was also greater in PLK1-quercitrin and PLK1-kaempferol 3-O-beta-D-glucoside, as the mutation led to energy losses of −5.17±0.02 kcal/mol and −2.64±0.02 kcal/mol, respectively. In the case of the PLK1-astragalin complex, the mutation of GLU131 to alanine also resulted in a notable decrease in the binding free energy, with a ΔΔ_TOTAL_ of −1.00 ± 0.06 kcal/mol. ASP194 and GLU131 appear to be involved in essential hydrogen bonding and electrostatic interactions, which are disrupted upon mutation to alanine, leading to a decrease in binding affinity. This disruption likely results from the loss of side chain functional groups that participate directly in binding interactions, emphasizing the role of specific amino acids in the molecular recognition process ^52^.

According to the analysis of PDK1 complexes, the mutation of LYS163 to alanine significantly impacts the binding affinity across three specific complexes: luteolin 7-O-(6-malonylglucoside), thermopsoside, and scolimoside. This mutation resulted in considerable energy losses of −15.11±0.05 kcal/mol, −5.67±0.03 kcal/mol, and −3.49±0.03 kcal/mol, respectively, indicating that LYS163 plays a critical role in stabilizing ligand□protein interactions. The negative ΔΔ_TOTAL_values underscore the importance of LYS163 in maintaining favourable electrostatic interactions and hydrogen bonding with ligands, which are significantly compromised upon mutation to alanine, a residue incapable of forming such specific interactions ^53^. Conversely, mutating ALA162 to glycine in the PDK1-scolimoside and PDK1-thermopsoside complexes yielded positive energy changes, with ΔΔ_TOTAL_ values of 12.49±0.38 kcal/mol and 9.55±0.31 kcal/mol, respectively. These positive shifts suggest that the glycine mutation of ALA162 has a relatively minor detrimental impact on the binding affinity ^54^.

In summary, this computational study provides compelling evidence for the use of bioactive compounds from Daucus carota as potent inhibitors of PLK1 and PDK1 for TNBC treatment. With favourable drug-like properties and multitargeted inhibition, these novel phytochemicals can pave the way for more efficacious and safer targeted therapies for aggressive TNBC tumors.

## 4. Conclusion

This study sheds light on the potential of Daucus carota-derived phytochemicals as novel therapeutic agents targeting PLK-1 and PDK-1 for triple-negative breast cancer (TNBC) treatment. Our in-silico investigation identified six promising lead compounds, namely, astragalin, kaempferol-3-O-β-D-glucoside, quercitrin, scolimoside, luteolin-7-O-(6-malonylglucoside), and thermopsoside, which exhibit favourable binding affinities and interactions with key residues in the ATP-binding sites of PLK-1 and PDK-1. Notably, compared with those of the established drug epirubicin, the binding energies of astragalin and scolimoside were superior. Molecular dynamics simulations further bolstered the confidence in these leads by demonstrating the remarkable stability of Astragalin and kaempferol-3-O-β-D-glucoside at the PLK-1 ATP site and of scolimoside within the PDK-1 binding pocket throughout a 150 ns simulation. Additionally, computational alanine scanning revealed critical residues, such as ASP194 and GLU131 for PLK-1 and LYS163, LYS169, and GLU166 for PDK-1, that play instrumental roles in ligand binding. Collectively, our findings illuminate a promising avenue for the development of plant-based PLK-1 and PDK-1 inhibitors, potentially paving the way for more effective and targeted therapeutic strategies against aggressive TNBC tumors. While further in vitro and in vivo experimentation is warranted to validate the anticancer efficacy and pharmacological properties of these lead compounds, this study establishes a robust framework for future investigations into harnessing the therapeutic potential of nature’s bounty for combating TNBC.

## Author contributions

Conceptualization: Kayode Raheem; Introduction: Modinat Abayomi, Mariyam Oluwatosin, Mary Adewunmi; Formal analysis: Kayode Raheem, Modinat Abayomi; Data curation: Kayode Raheem. Writing original draft preparation: Kayode Raheem, Modinat Abayomi, and Mariyam Oluwatosin; Writing review and editing: Kayode Raheem, Modinat Abayomi, and Muhammad Muddassar. Supervision: Muhammad Muddassar. All the authors have read and approved the final draft.

## Acknowledgements

We are grateful to the Chemical Biology and Drug Designing Lab under the supervision of Dr. Muhammad Muddassar for providing hardware and software support. In addition, we would like to acknowledge the support we received from the Cancer Research with AI (CARESAI) group at the completion of this project.

